# N-terminus of the third PDZ Domain of PSD-95 Orchestrates Allosteric Communication for Selective Ligand Binding

**DOI:** 10.1101/2020.08.24.264226

**Authors:** Tandac F. Guclu, Nazli Kocatug, Ali Rana Atilgan, Canan Atilgan

## Abstract

PDZ domains constitute common models to study single domain allostery without significant structural changes. The third PDZ domain of PSD-95 (PDZ3) is known to have selective structural features that confer unique modulatory roles to this unit. In this model system two residues, H372 directly connected to the binding site and G330 holding an off-binding-site position, were designated to assess the effect of mutations on binding selectivity. It has been observed that the H372A and G330T/H372A mutations change ligand preferences from class I (T/S amino acid at position −2 of the ligand) to class II (hydrophobic amino acid at the same position). Alternatively, the G330T single mutation leads to the recognition of both ligand classes. We have performed a series of molecular dynamics (MD) simulations for wild-type, H372A, G330T single mutants and a double mutant of PDZ3 in the absence and presence of both types of ligands. With the combination of free energy difference calculations and a detailed analysis of MD trajectories, ‘class switching’ and ‘class bridging’ behavior of PDZ3 mutants, as well as their effects on ligand selection and binding affinities are explained. We show that the dynamics of the charged N-terminus plays a fundamental role in determining the binding preferences in PDZ3 by altering the electrostatic energy. These findings are corroborated by simulations on N-termini truncated versions of these systems. The dynamical allostery orchestrated by the N-terminus offers a fresh perspective to the study of communication pathways in proteins.

## Introduction

PDZ domain-containing protein complexes control intercellular interactions such as trafficking, signaling, cell to cell communication and organization of signaling complexes (1–3). A wide variety of organisms from bacteria to vertebrates contain varying numbers of PDZ domains (3, 4). As one of the PDZ containing protein complexes, PSD-95 has three PDZ domains followed by the Src Homology 3 (SH3) and guanylate kinase (GK) domains (5). The role of each PDZ domain in PSD-95 as well as how they function in tandem is actively researched (see e.g. ref. (6)).

PDZ domains are themselves small proteins which typically consist of 90 to 100 amino acids and have 6-7 *β*-strands and two *α*-helical structures as characteristic features (1–3, 5, 7). Common structural motifs are consistent in the folding respect, but differ in length amongst the PDZ domain family (8). In addition to its secondary structure, the loop at the binding cleft, referred to as the carboxylate-binding loop, is one of the hallmarks of the PDZ domain (9). Members of the PDZ domain family bind selectively to short amino acid patterns at the C-termini of targeted proteins (3, 8). Therefore, the pattern of the target protein is the definitive factor for the classification of PDZ domains (1, 10). PDZ domain-binding protein interactions are categorized based on the amino acid at the second position of the binding protein downstream from the C-terminus (9, 11). Ligands are classified as Class-I (CRIPT) (12) for Thr/Ser, Class-II for hydrophobic amino acids and Class-III for Asp/Glu (9, 10, 13, 14) at this position. On the other hand, there is a selective structural feature of the third PDZ domain of PSD-95 (PDZ3) which makes it more interesting to work on; it has an extra α helical structure at the C-terminus (15–17). Experimental and computational studies reveal the significance of the unusual α3-helix as a stabilizing effect and its participation in the “hidden” allosteric communication (4, 15–20). Furthermore, PDZ3 was previously utilized to demonstrate that dynamical allostery is not only due to entropic factors, but also might involve redistribution in energetics, particularly of electrostatics origin, accounting for population shifts observed in dynamical allostery (21, 22).

In this study, PDZ3 is used as a model protein. For this model, Class I (L_1_) and Class II (L_2_) ligand-binding are possible (Figure 1). Two residues have been found crucial in affecting the ligand preference; G330 and H372 (23–25). G330 does not have a direct contact with the ligands, and is located on a nearby loop (23). Conversely, H372 is at the binding pocket of the protein and is crucial for L_1_ binding (16). It has been shown that mutations on these two residues alter the affinity of the protein to different classes of ligands (23, 25). The wild type (WT) prefers to bind L_1_ (Figure 1a); however, once G330T mutation occurs, the PDZ domain has similar affinity to both L_1_ and L_2_. Due to the resultant change in ligand preference, G330T-type mutations are called *class bridging* (23–25) (Figure 1b). H372A mutation has a decreasing effect on L_1_ binding affinity, while simultaneously increasing affinity for L_2_ (23–25). Therefore, mutations such as H372A are called *class changing* mutations (23) (Figure 1c). If the binding protein undergoes both G330T and H372A mutations (termed double mutant, DM, in this study), the protein binds L_2_ only; i.e. the double mutation is also class changing (23) (Figure 1d). While these observations have been studied extensively both experimentally (14, 23, 25–29) and computationally (24, 30–32), the governing mechanisms are not clear as of yet; moreover, most of these studies address structural features induced by these changes while a systematic dynamical assessment is not available.

**Figure 1.**
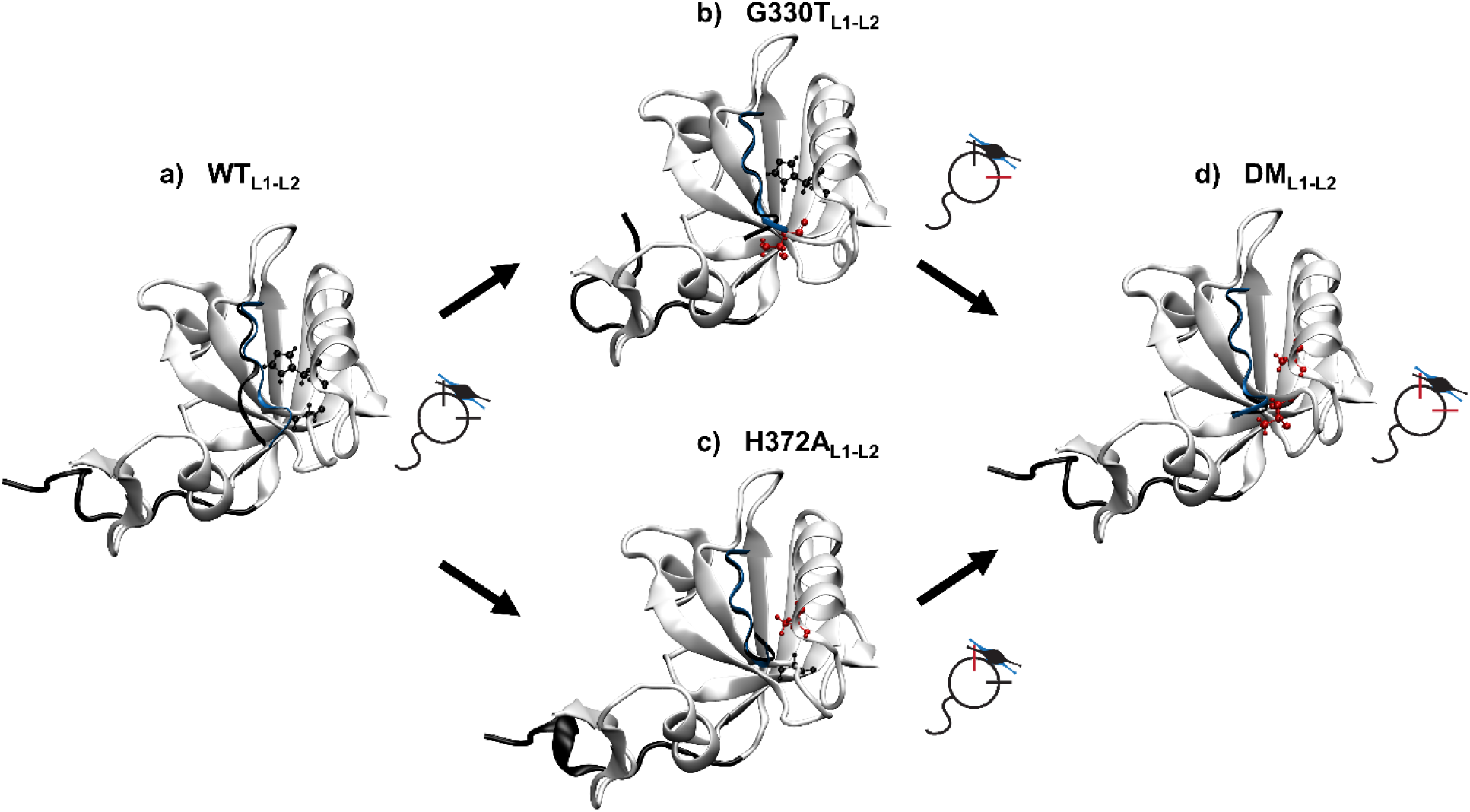
PDZ domain complexes for WT, G330T, H372A and DM cases are visualized with the mutations and the ligands (PDB codes listed in Table 1). On the protein structure, G330 and H372 sites are shown in black for the wild-type form, and the mutated residues are shown in red. CRIPT (L_1_) and N-terminus region (residues between 299 and 310) are shown as black ribbons, while T-2F (L_2_) is illustrated in blue. The shorthand cartoons are used to differentiate each case: Circle represents the body of the protein; ticks, whiskers and violin shapes correspond to the mutation sites, the N-terminus regions and the ligands, respectively. The color code for each component in cartoons is the same as the protein structure. Residue 330 is not directly located at the binding site, whereas 372 directly interacts with the ligand. **a** WT structure binds strongly to the L_1_. **b** G330T mutant is the class bridging mutant; thus, it binds both L_1_ and L_2_ with similar free energy difference. **c, d** H372A and DM structures prefer to bind L_2_ with a lower energy value; therefore, H372A mutation is defined as the class switching mutant.

We perform molecular dynamics (MD) simulations for the apo and L_1_/L_2_-bound form of the WT, G330T, H372A and DM PDZ3 structures. By using the conformations obtained from the MD trajectories, we conduct free energy perturbation (FEP) simulations (33, 34) to investigate the energy cost of the mutations. Integrating the binding and mutation energies into thermodynamic cycles allows us to relate the computational results to the experimental binding energies from a previous study (23). Further, our detailed analyses of the MD trajectories reveal that the charged N-terminus region of PDZ3 has a significant impact on the ligand specificity in addition to the direct interactions occurring at the binding site. To test this hypothesis, we replicate the simulations on the N-terminus truncated complexes. We show that the free energy differences leading to the class bridging/switching behavior are nullified in the absence of the N-terminus. Thus, we demonstrate the electrostatic contributions due to the dynamics of the N-terminus region is essential for the formation of the functional PDZ complexes.

## Results

### Relative binding energies select functional ligands for different mutations

Thermodynamic cycles to compute various relative free energy changes due to mutations or ligand binding are constructed as illustrated in Figure 2a. For each cycle connected by solid arrows, the difference between two vertical free energy changes (Figure 2a, red) signifies the more stable bound form; i.e. for a negative ΔΔ*G*_A_ (see Equation 4 in Methods) the mutated complex with L_1_ is more favorable than the WT complex. Since each of the cycles completed by the solid arrows should sum to 0, the ΔΔ*G* calculated from experimental binding constants may also be obtained by computational means, from the difference between the two horizontal free energy changes (Equations 5–6).

**Figure 2.**
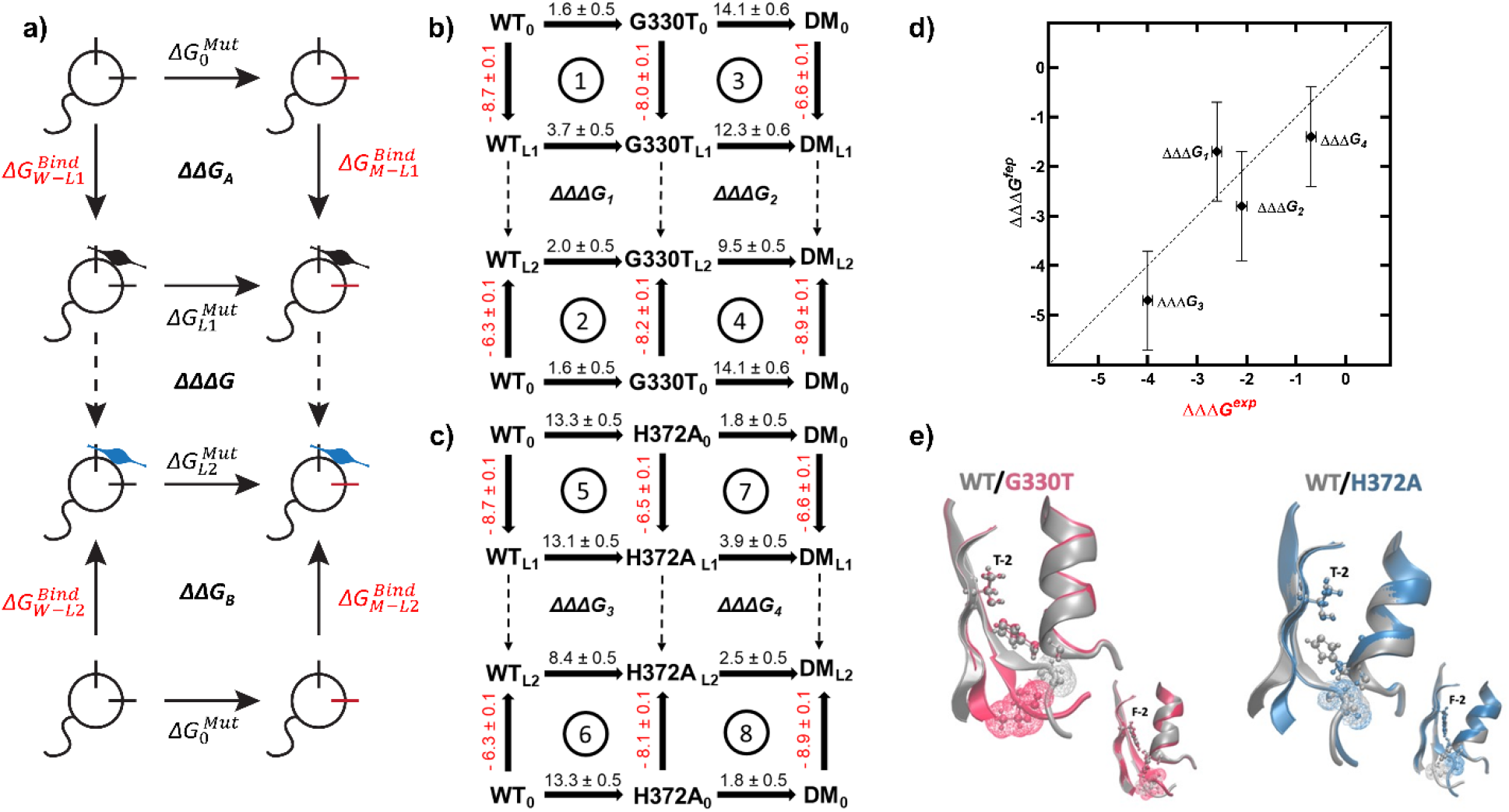
**a** Cartoon illustration for mutation and ligand binding thermodynamic cycles of the PDZ domain. Violin shapes in black and blue represent L_1_ and L_2_, respectively. Δ*G*^Mut^ values (in black) show the FEP simulation derived free energy differences between the WT and mutated PDZ3. Δ*G*^Bind^ values (in red) are the standard binding free energy differences calculated from experimentally reported *K_d_* (23). Dashed arrows indicate the ligand switching process (not directly calculated). **b, c** Mutational cost (black) and ligand binding free energies (in red) for G330T, H372A and DM; all values in kcal/mol. ΔΔΔ*G* may be calculated by using either Δ*G*^Mut^ or Δ*G*^Bind^ values. **d** Plot of FEP calculated vs. experimental ΔΔΔ*G* (Pearson *R* = 0.84, *p*-value < 0.01) attest to the range of validity of FEP calculations. **e** Close-up views of side chain positioning and residue 330 solvent accessibility affected by the mutations (WT gray, G330T pink, H372A blue). L_1_ bound forms are displayed large, L_2_ bound forms are small. G330T mutation (left) does not affect interactions between H372 and position −2 in either ligand but leads to a conformation shift in the loop carrying position 330, increasing its solvent accessibility. H372A mutation (right) in L_1_ bound form leads to loss of polar-polar interactions in the binding pocket; in L_2_ bound form, the kink in the loop containing G330 due to rotation of H372 side chain to accommodate the large F-2 is relieved upon mutation. Solvent accessibility of G330 remains the same in both cases.

The binding constants from a previous study (23) are employed to calculate the standard binding free energies of WT, G330T, H372A and DM PDZ3 to L_1_ and L_2_ (shown in red in Figures 2b-c). Since PDZ domains were shown to operate at the 1-15 μM dissociation constant range (35), following previous work we have used −7.0 kcal/mol as a standard binding free energy threshold (~10 μM at physiological temperatures). Therefore, WT_L1_, G330T_L1_, G330T_L2_, H372A_L2_ and DM_L2_ are the functional PDZ complexes based on their binding free energies, which are −8.7, −8.0, −8.2, −8.1 and −8.9 kcal/mol, respectively. The binding free energies of non-functional complexes are −6.3, −6.5 and −6.6 kcal/mol for WT_L2_, H372A_L1_ and DM_L1_, respectively. The FEP calculated free energy costs of the mutations are also displayed in the same figures in black.

Note that the tautomeric states of key residues, such as H372, may change upon ligand binding and that they may have significant populations in multiple states in bound or apo forms (36, 37). Since each cycle should sum to zero, we find from the deviations in cycles ①, ②, ⑦ and ⑧ that the apo form G330T mutation has an additional cost of ~1.8 kcal/mol, and from those in cycles ③, ④, ⑤ and ⑥ that the H372A mutation in the apo form has an additional ~3.2 kcal/mol cost due to such population shifts.

Although each leg of the cycles in Figures 2b-c may be obtained computationally, such calculations are subject to various errors inherent in the employed methodology. In Figure 2a, ΔΔΔ*G*, i.e., the difference between the dashed arrows is equal to ΔΔ*G*_B_ – ΔΔ*G*_A_ (horizontal) as well as that between the vertical binding free energies (Equation 7). Thus, we employ ΔΔΔ*G* to validate our simulations against the experimental binding energies (Figure 2d) and we find that within the 1 kcal/mol accuracy limit provided by FEP calculations (38), the energy differences obtained by using conformations sampled throughout the MD trajectories are consistent with the experimental binding energies. Note in particular that this approach eliminates the use of mutational cost calculations in the apo forms, in particular the above-mentioned energy cost due to shifts in dynamic tautomeric populations.

Since we do not calculate binding free energies, we are not positioned to directly comment on which mutants will be functional. However, we are now able to discuss the mechanisms by which these mutations operate. First off, the general effect of ligand type on free energetic cost of the mutations is clear from the calculations where changes in the L_2_ bound form is always less costly than those in the L_1_ bound form, hence the negative ΔΔΔ*G* values displayed in Figure 2d.

As discussed in a previous study on the mutational preferences of PDZ (23), G330T is the most abundant variant owing to its class bridging behavior (Figure 2e, left). This phenomenon is explained by the low free energy cost of this mutation, irrespective of the presence of the ligand in the binding pocket and of the prior H372A mutation (in the range of up to 4 kcal/mol; Figure 2, cycles ①, ②, ⑦ and ⑧; numbers in black). Considering that the G330T mutation requires relatively low number of atom additions and that residue 330 is located on a flexible loop (Figure 2e, left) where the newly created polar side chain may easily be repositioned to get in contact with water, the low cost is plausible. Nevertheless, the L_2_ bound form is able to accommodate this mutation with ~2 kcal/mol less cost than either the ligand free or the L_1_ bound forms, leading to the ligand bridging behavior (compare cycles ① and ②). Similarly, the H372A mutant is also able to contain the additional G330T mutant with ~1.5 kcal/mol less cost in the presence of L_2_ (compare cycles ⑦ and ⑧), retaining the ligand switching behavior brought on by this first mutation.

In contrast to G330T, the H372A mutation is rather costly (on the order of 10 kcal/mol) under all circumstances due to the large change of the side chain volume, as well as the shift this position makes from polar to hydrophobic (Figure 2e, right). Moreover, this residue is in the binding pocket in direct contact with the ligand in the liganded cases, which makes the free energetic cost highly dependent on the rearrangements of local contacts. Especially in L_1_-bound cases, the bond between H372 and the threonine residue at the −2 position of L_1_ is pertinent to the formation of the WT_L1_ complex (16). Note that H372 was also shown via deep mutagenesis to be the most sensitive residue to mutations (25). Nevertheless, the cost of the H372A mutation is significantly lower when L_2_ is bound to the protein, by 4.7 kcal/mol (compare cycles ⑤ and ⑥), helping bring the binding free energy of this mutant in the functional range, and thus leading to ligand switching. When H372A mutation follows G330T, the cost is again lower in the presence of L_2_, but more importantly, it is enough to compensate for the effect of the mutation in the apo form and to retain a physiologically significant degree of specificity.

### N-terminus fluctuation patterns are distinct for each bound ligand and mutant type

The ligand-bound trajectories we have generated reliably represent the equilibrium properties of the bound complexes, as validated in the previous subsection since the FEP calculations are based on the conformations generated in these simulations. We now focus on the dynamics of the complexes to delineate the allosteric behavior observed in these systems. We thus compute the overall root mean square deviation (RMSD) of each PDZ3 from the initial x-ray structure, the root mean square fluctuations (RMSF) and the cross-correlations of the residues averaged over the trajectories.

RMSD for each MD trajectory is shown in Figure 3. We find an unusual property for these trajectories whereby there are stretches of times having plateaus with small fluctuations, separated by regions of relatively large fluctuations. RMSF results show that the most mobile site of PDZ3 is the N-terminus region (Figure S1). Hence, we determine that the overall mobility of the protein is dominated by the fluctuation of the N-terminal region. All apo complexes have highly mobile and disorganized N-termini, while those of the ligand-bound complexes have clusters around preferred conformational states (Figure 3). The ligand-bound forms of WT and G330T have similar cluster shapes for the N-terminus residues (Figure 3a-b). H372A_L1_ and DM_L1_ exhibit nearly identical conformational preferences of the N-terminus (Figure 3c-d). The N-terminus of H372A_L2_ adopts an extended conformation, while that of DM_L2_ displays a collapsed form (Figure 3c-d).

**Figure 3.**
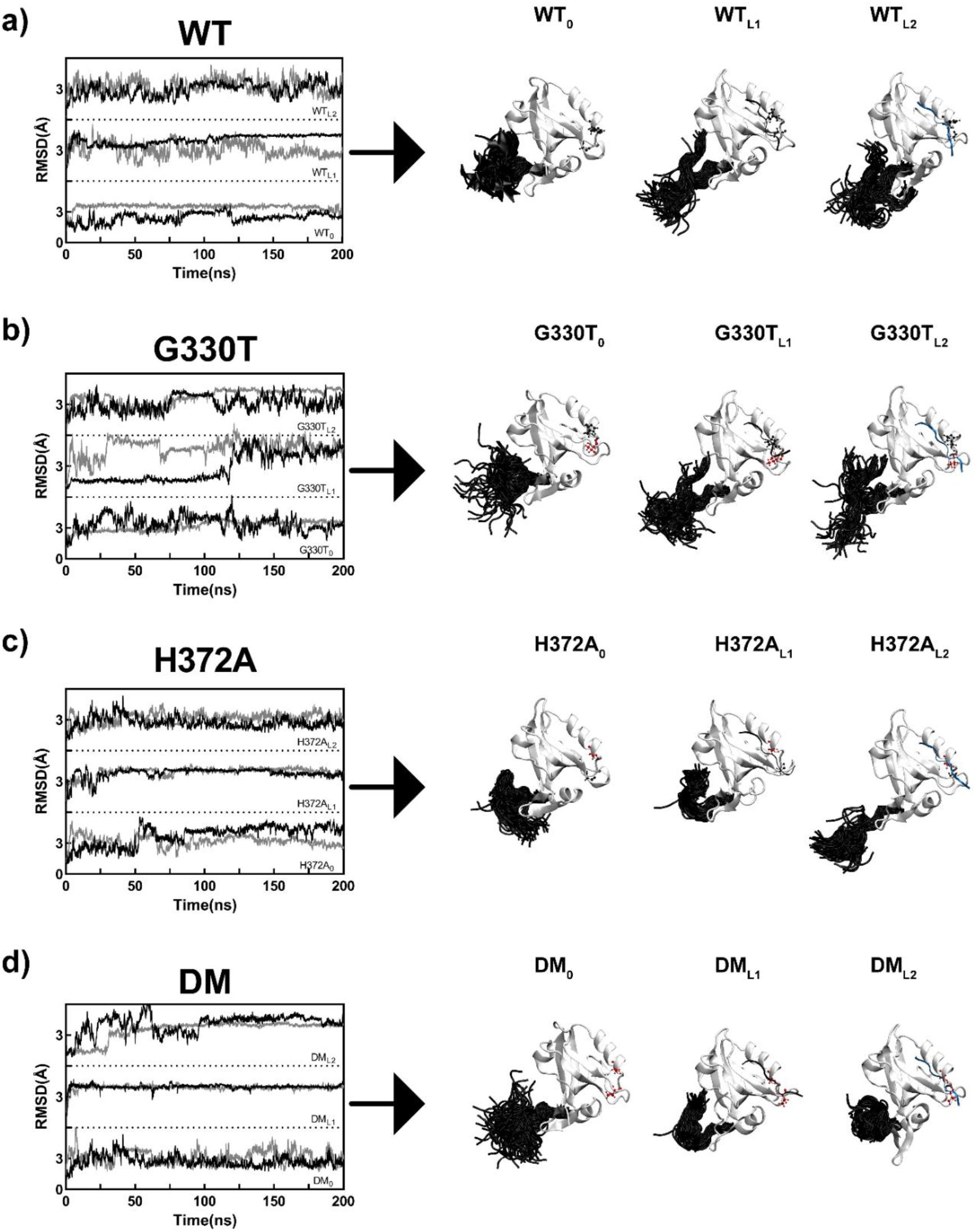
RMSD and N-terminus conformers observed in the MD simulations for apo and L_1_/L_2_ bound cases. In RMSD plots, duplicate simulations are displayed separately by black and gray curves. In protein structure visualizations, evenly separated 500 snapshots of only the N terminus region (shown in black) from duplicate trajectories for each complex are illustrated; the rest of the protein (in white) is displayed by its average structure with side chains of residues 330 and 372 displayed in ball and stick (red if mutated, black if native). L_1_ and L_2_ are shown in black and blue, respectively.

To scrutinize the dynamical behavior of the PDZ structures further, the cross-correlation matrices of the protein complexes are calculated (Figure S2). The correlations of residue displacements are very similar, with the inner product between WT_0_ and each of the other systems varying in the range 0.80-0.97 when the 12 residue-long N-terminus spanning residues 299-310 is omitted in the calculations. Thus, the cross correlations of the protein are dominated by the ordered main body while the N-terminus region adopts unique dynamics depending on the adopted point mutations and bound ligands.

### N-terminus is an allosteric partner essential in determining binding ligand

The N-terminus has the NH_2_-P**E**FLG**EED**IP**RE** sequence; half of its 12 residues are charged, and the rest are hydrophobic. It also renders the −4 net charge of the protein. This sequence possibly contributes to the long-range control over binding affinities. To quantify the various degrees of flexibility observed in the conformations discussed in the previous section (Figures 3, S1 and S2), we plot the solvent accessible surface area (SASA) distributions of the N-terminus residues (Figure S3). The disordered flexibility of the region in the apo forms is characterized by their broad distributions, while SASA also delineates the two-state nature of the conformations of the WT_L1_, G330T_L2_ and even distinguishes minor conformations such as that observed for G330T_L1_; DM_L2_ is particularly characterized by a peakish single conformation. SASA distributions of only the charged N-terminus residues, on the other hand, display starkly different features (Figure 4). First, those of the apo forms all have the same type of broad distribution, with variance ~60 Å^2^. More interestingly, while the whole N-terminus may take alternate conformations, it does so at the expense of confining the solvent exposure of its charged residues at a given value; this is best exemplified by WT_L1_. Note that the average number of hydrogen bonds established between PDZ3 and the ligands (varying between 3 and 4) also depend on the mutant type (displayed in Figure 4 sidebars and listed in Table S1). Although, the difference is small, hydrogen bonds established between protein and ligand have a significant impact on ligand binding (39). Moreover, there is a meaningful negative correlation between the average SASA of the N-terminus and the number of hydrogen bonds (Pearson *R* = −0.86; Figure S4a) implying allosteric communication between binding site and the charged tail.

**Figure 4.**
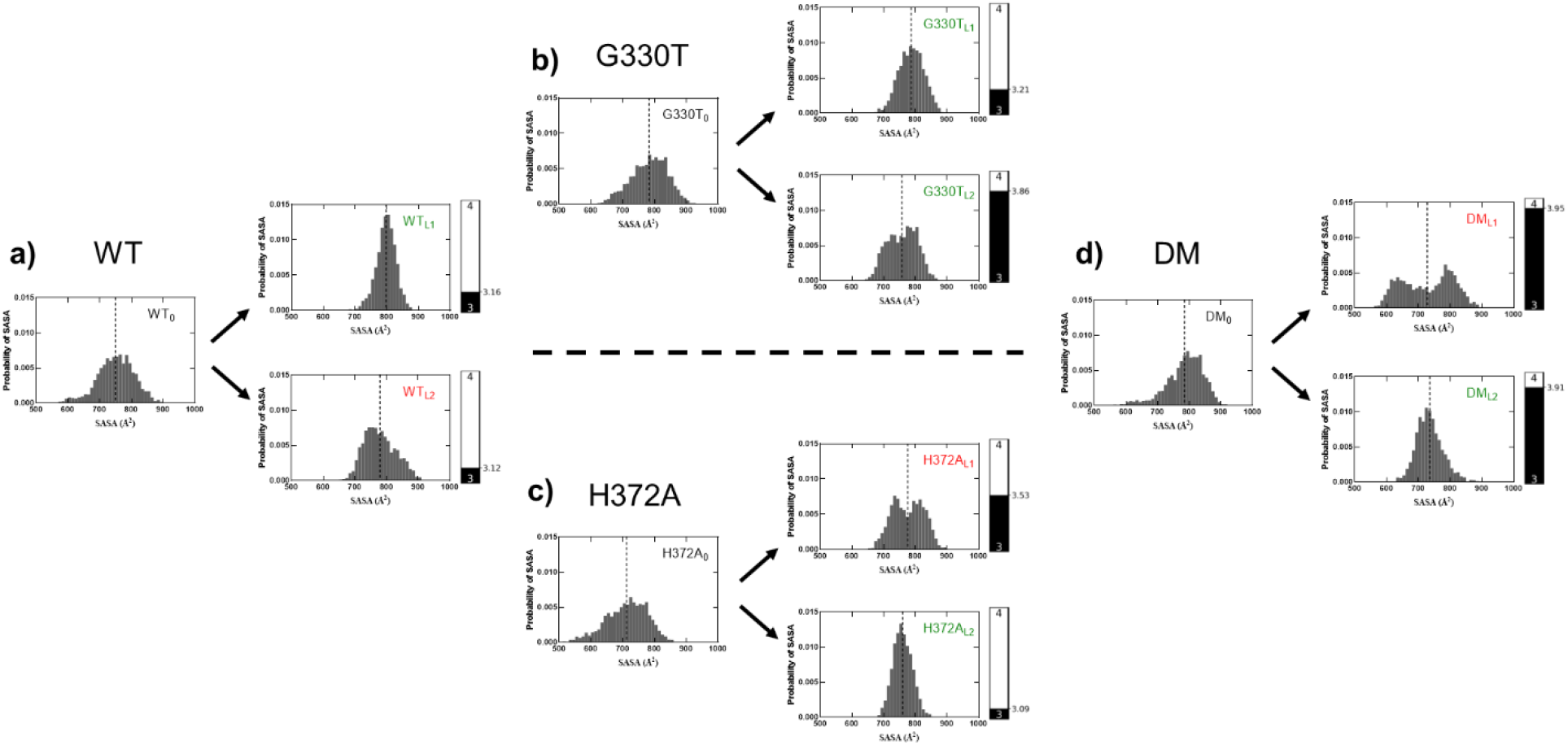
Probability distribution of SASA for the N-terminus charged residues; side bars display the average hydrogen bond count between the PDZ domain and the ligand in the MD simulations in ligand-bound cases. The complexes with favorable binding to the ligands are labelled in green, while the complexes with unfavorable binding are in red. **a** Regarding the WT complexes, solvent accessibility of the charged residues has a dominant effect on ligand-binding. WT_0_ and WT_L2_ have similar broad distributions while WT_L1_ is peakish leading to favorable binding. Average hydrogen bond counts are indifferent in both cases. **b** SASA profiles of G330T complex do not differ drastically after binding. The L_1_ bound complex has a slightly sharper peak and a lower hydrogen bond count. However, the L_2_ bound form has a wider SASA distribution and a higher hydrogen bond count. Additional hydrogen bonds and sharper SASA distributions compensate one another to induce the favorable binding in the alternate cases. **c, d** In H372A and DM cases, the favorable binding to L_2_ is mainly due to the sharp SASA distribution of the charged residues on the N-terminus.

To explain the observed binding preferences of PDZ, we have identified two main contributions: (i) The direct effect at the binding site quantified by the average number of hydrogen bonds between the protein and the ligand; and (ii) the allosteric effect due to conformational multiplicity of the N-terminus and its resulting dynamic interactions. The latter is related to the electrostatic free energy change of the system (40, 41),

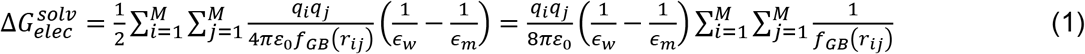

While not defined uniquely, several simple but effective forms were used for the function *f_GB_*(*r_ij_*) including inverse relation to charge-charge distances *r_ij_* and screening effects. Here, we assume 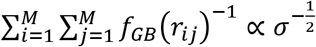, where *σ* is the variance of the SASA of the charged residues. Moreover, the two contributions also determine the overall conformations adopted by the protein. Thus, our simplified model to predict binding free energies from MD trajectories is given by the sum of three effects:

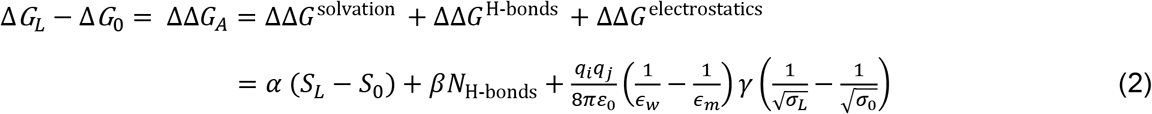

The first is the usual term accounting for the change in the solvation free energy of the protein (42–46), proportional to the difference in the average SASA of the whole protein in the ligand-bound and ligand-free forms, *S_L_* and *S*_0_, respectively (Table S2). The second term accounts for the interaction energy between the ligand and the binding site, dominated by the number of hydrogen bonds formed at the binding cavity for each ligand-bound trajectory, *N_H-bonds_*. The last term approximates the electrostatic free energy change, with *ε_w_* and *ε_m_* being the dielectric constants of water and the buried medium, respectively, and *σ_L_* and *σ*_0_ representing the variance in the SASA of the charged residues of the N-terminus for the liganded and ligand free forms, respectively (Table S2). Equation 1 regresses the experimental free energy of binding data with *α* = 2 cal/mol/Å^2^, *β* = −1.6 kcal/mol, and *γ* = 0.75 for *ε_w_* = 80 and *ε_m_* = 2 (Pearson *R* = 0.94; Figure S4b). These pre-factors are physiologically relevant: Previously reported data for *α* is in the range of a few to tens of cal/mol/Å^2^ (43, 47–49), and for *β*, the rupture of each hydrogen bond may cost the total energy of a protein ~1.6 kcal/mol in aqueous environment (50–52).

The contribution of each term to the binding free energy is listed in Table 2. Note that the largest contribution is due to the hydrogen bonds established in the binding pocket in each case. Nevertheless, the overall conformational change of the protein as well as the specific conformations adopted by the N-terminus charged residues determine the binding fate.

Thus, the L_1_ preference of the WT protein is reinforced by the fixed conformations adopted by the N-terminus charged residues, despite similar hydrogen bonding interactions it has with both ligands (Table 2 and Figure 4). The G330T single mutation is a special case which prefers to bind both L_1_ and L_2_. However, the preference to L_1_ is reinforced by the electrostatic contributions of the complex, while that to L_2_ is due to the increased number of hydrogen bonds established at the binding cavity. Accordingly, the ligand-bridging behavior of the G330T mutation is conducted by the compensation between the conformational dynamics of the N-terminus region and the protein-ligand hydrogen bonding. The H372A single mutation prefers to bind L_2_ due to significant contributions from the N-terminus electrostatics, a factor that is nearly absent for the L_1_ bound complex. In the presence of both mutations, we find that there is significant increase in binding pocket interactions as well as the overall solvation free energy of both DM_L1_ and DM_L2_ complexes. However, the N-terminus charged residue fluctuations in DM_L1_, which uniquely exceed those in the unbound form (with a positive ΔΔ*G*^electrostatics^) offsets this advantage for L_1_.

### Removal of the charged N-terminus exposes its key role in ligand binding

We design a knock-out computer experiment by removing residues 299-310 constituting the N-terminus to test its role on binding affinities; in what follows, these systems are referred by the superscript Δ. We run 50 ns-long MD simulations for the N-terminus deleted forms of all eight ligand bound systems. Additionally, we perform FEP calculations for the single mutations (G330T^Δ^ and H372A^Δ^).

In Figure S5a, we display the free energetic cost of removal the N-terminus from the two single mutations. In the full-length PDZ, the N-terminus appears to have the largest favorable impact on WT_L1_, G330T_L1_ and H372A_L2_ (Table 2). Thus, we expect cycle ② to be the least affected by its absence as is the case, with the G→T transition in the presence of L_2_ costing 2.0 ± 0.5 to 1.9 ± 0.3 kcal/mol, for G330T and G330T^Δ^, respectively. In cycles ①, ③ and ④ at least one of the constituents of the full-length proteins is highly dependent on N-terminus dynamics, and it is therefore not straightforward to judge which will cost the most. We do find through FEP calculations, however, that cycle ① which has both WT_L1_, G330T_L1_ involved is the most affected, with the cost of the G→T transition in the presence of L_1_ costing 3.3 kcal/mol more when the charged N-terminus is removed. This is followed by cycle ④ where the binding pocket H→A transition in the presence of L_2_ has an additional cost of 2.6 kcal/mol.

The loss of the N-terminus translates into modified binding pocket interactions where we may trace the origin of the differing free energy differences. Removal of N-terminus increases *N_H-bonds_* in all cases by an average of 0.2-0.5, except in 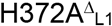 where it decreases by 0.5 (Table S1). For the G330T transition, L_2_ binding is more favorable, mainly because the deletion of the N-terminus leads to a net gain of nearly one full hydrogen bond (from 3.4 to 4.3) while it is a mere 0.3 gain (from 3.4 to 3.7) in its L_1_ bound form, also corroborated by 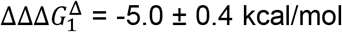 (Figure S5 b). For the H372A transition, L_2_ binding is again more favorable, but with a lower propensity 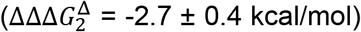, as the L_1_ and L_2_ binding cost −0.4, and +0.1 hydrogen bonds, respectively. Finally, we expect the DM^Δ^ systems to favor binding both L_1_ and L_2_ due to the gain of a full hydrogen bond in the binding pocket in the absence of the N-terminus. We thus expect that if the total binding free energy is in the functional range, then the ligand bridging behavior of the G330T^Δ^ mutation will change to ligand switching while the reverse is expected to happen for DM^Δ^.

## Conclusions

PDZ domains are parts of signaling proteins which play key roles in cellular functions (1–3). Here, we study the PDZ3 complex with a focus on the G330T and H372A mutants which bind to CRIPT (L_1_) and T-2F (L_2_) (16, 23–25). Experimental studies have shown that WT_L1_, G330T_L1_, G330T_L2_, H372A_L2_ and DM_L2_ are the functional complexes while others are unfavorable based on their binding constants (23, 25, 35). In this study, we perform MD and FEP simulations to show the energetic, conformational and dynamical bases for the ligand specificity of the mutations in PDZ domains.

The FEP simulations demonstrate the energy cost of individual mutations. The computational ΔΔG values are in agreement with the experimentally determined adaptive pathway, and the *in silico* ΔΔΔG results corroborate the experiments (23). The multiple MD simulations uncover intriguing fluctuation patterns of the glutamic acid rich N-terminus region (Figure 3) which provide the −4 net charge of the protein. Accordingly, to predict binding free energies from MD trajectories, the proposed model (equation 2) accounts for this electrostatic effect along with the change in the hydrogen bond contributions due to different mutations. Our model fits well the results of binding experiments; furthermore, its regression parameters are in the biophysically relevant range.

Our simple model demonstrates that the charged residues of the N-terminus have a decisive role in mutation – binding partner matches for functional activity, even though the main contributions to binding free energy come from the hydrogen bonds formed in the binding pocket (Table 2). Namely, (i) the H372A mutant prefers L_2_ because of the Born solvation energy of the N-terminus, which is negligible for the L_1_ bound complex; and (ii) while the G330T prefers to bind both L_1_ and L_2_, the preference to L_1_ is strengthened by the electrostatic contributions of the complex, similar to the choice of L_1_ in lieu of L_2_ for the wild type.

These observations lead to the following fundamental question: Would mutations select the same ligands functionally, if we were to intervene with the proposed communication between the N-terminus and the rest of the protein? To answer, we simply remove the 12 N-terminus residues from all of the ligand bound proteins, repeat the calculations (Figure S5), and monitor the changes in the binding energies. We find that the costs of the G→T transition in the presence of L_1_ in cycle ① and the binding pocket H→A transition in the presence of L_2_ in cycle ④ are 3.3 and 2.6 kcal/mol larger, respectively. This finding clearly summarizes the moderating role of the N-terminus for selecting the functional ligand for the PDZ domain. Our further analysis via the proposed simple binding free energy model indicates that the FEP differences in the truncated PDZ3 are mainly due to modified hydrogen bond interactions in the binding pocket.

To sum up, our results demonstrate that the unstructured N-terminus region is fundamental for the so-called “hidden dynamic allostery” in PDZ domains (53). In particular, it unravels the origins of the electrostatic basis leading to the dramatic population shifts observed in the pairwise hydrogen-bonded interactions involving the side chains (22). Future studies linking the N-terminus fluctuations to stabilizing α3-helix will provide a more complete picture of the mechanisms leading to regulation of binding partners by PDZ domains when performing their signaling roles in the cell.

## Methods

### MD simulations

PDZ3 structures are obtained from the PDB (54) (Table 1). Where necessary, the protein lengths are equated by deleting residues 297 and 298 from the N-terminal, and modeling missing residues of the ligands by using SWISS-MODEL server (55). MD simulations are carried out with the NAMD (56) package under the CHARMM36 (57) force field. The simulations are visualized using VMD (58). Protein structures are placed into a water box whereby a minimum distance of 10 Å is maintained between each atom of the protein and the nearest edge of the water box by using the solvent plug-in VMD 1.9.3. Residue protonation states are consistent with pH 7.4. By adding a sufficient number of potassium chloride ions (KCl) to the system, the ionic strength is adjusted to 150 mM while achieving charge neutrality. Long-range electrostatics are calculated by the particle mesh Ewald method (59), with a cut-off distance of 12 Å. Temperature control is maintained by Langevin dynamics. The system is run under 1 atm and 310 K in the NPT ensemble. The complete system is minimized for 10,000 steps and equilibrated for 100,000,000 steps; the time step is 2 fs.

**Table 1.** Model proteins and their PDB IDs.

| **Abbreviation** | **PDB ID** | **Structure Details** |
| --- | --- | --- |
| WT<sub>0</sub> | 5HDY | PDZ<sup>3</sup> of PSD-95 |
| G330T<sub>0</sub> | 5HET | PDZ<sup>3</sup> of PSD-95 (G330T mutant) |
| H372A<sub>0</sub> | 5HF4 | PDZ<sup>3</sup> of PSD-95 (H372A mutant) |
| DM<sub>0</sub> | 5HFD | PDZ<sup>3</sup> of PSD-95 (G330T, H372A double mutant) |
| WT<sub>L1</sub> | 5HEB | PDZ<sup>3</sup> of PSD-95 in complex with the peptide derived from CRIPT |
| G330T<sub>L1</sub> | 5HEY | PDZ<sup>3</sup> of PSD-95 (G330T mutant) in complex with the peptide derived from CRIPT |
| H372A<sub>L1</sub> | 5HFB | PDZ<sup>3</sup> of PSD-95 (H372A mutant) in complex with the peptide derived from CRIPT |
| DM<sub>L1</sub> | 5HFE | PDZ<sup>3</sup> of PSD-95 (G330T, H372A double mutant) in complex with the peptide derived from CRIPT |
| WT<sub>L2</sub> | 5HED | PDZ<sup>3</sup> of PSD-95 in complex with the mutant peptide derived from CRIPT (T-2F) |
| G330T<sub>L2</sub> | 5HF1 | PDZ<sup>3</sup> of PSD-95 (G330T mutant) in complex with the mutant peptide derived from CRIPT (T-2F) |
| H372A<sub>L2</sub> | 5HFC | PDZ<sup>3</sup> of PSD-95 (H372A mutant) in complex with the mutant peptide derived from CRIPT (T-2F) |
| DM<sub>L2</sub> | 5HFF | PDZ<sup>3</sup> of PSD-95 (G330T, H372A double mutant) in complex with the mutant peptide derived from CRIPT (T-2F) |

**Table 2.** Modeled vs. measured binding free energies and individual contributions from equation 2 (kcal/mol).*

| Complex | $\Delta\Delta G^{\text{solvation}}$ | $\Delta\Delta G^{\text{H-bonds}}$ | $\Delta\Delta G^{\text{electrostatics}}$ | Modelled $\Delta G^{\text{Bind}}$ | Measured $\Delta G^{\text{Bind}}$ |
| --- | --- | --- | --- | --- | --- |
| WT<sub>L1</sub> | -0.76 | -5.05 | **-3.00** | -8.81 | -8.65 |
| WT<sub>L2</sub> | -0.78 | -4.99 | -0.71 | -6.49 | -6.30 |
| G330T<sub>L1</sub> | -0.59 | -5.13 | **-1.82** | -7.54 | -8.02 |
| G330T<sub>L2</sub> | -0.71 | **-6.18** | -1.10 | -7.99 | -8.15 |
| H372A<sub>L1</sub> | -0.74 | -5.65 | -0.82 | -7.21 | -6.48 |
| H372A<sub>L2</sub> | -0.34 | -4.94 | **-3.19** | -8.47 | -8.11 |
| DM<sub>L1</sub> | **-1.53** | **-6.31** | **0.97** | -6.88 | -6.60 |
| DM<sub>L2</sub> | **-1.59** | **-6.25** | -1.44 | -9.28 | -8.94 |

\* functional complexes are displayed in green; interactions dominating the binding fate are shown in bold.

The 12 systems listed in Table 1 are simulated for duplicate 200 ns runs, resulting in 24 trajectories for a total of 4.8 μs simulations. In addition, the N-termini deleted versions of the ligand-bound forms are simulated for 50 ns each.

### Trajectory analyses

In RMSF, cross-correlation, hydrogen-bond occupancy and SASA calculations, the first 80 ns of the MD simulations are excluded. The remaining 120 ns is divided into three equal 40 ns chunks. The first frame of each trajectory is used as a reference for RMSD and RMSF calculations. To observe the correlated motions between residues, *N*×*N* crosscorrelation matrix is calculated from the trace of each 3×3 element of the covariance matrix (60). For visualization purposes, cross-correlation matrices are normalized between −1 and 1. Hydrogen bond and SASA analyses are performed in VMD with the default settings (58).

### FEP calculations and energy cycles

The free energy difference is calculated between reference (A) and target (B) states through the coupling parameter, λ (0→1) (61). While calculating free energy changes, the system space is sampled in forward and backward directions. The free energy difference is calculated via (62),

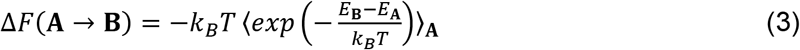

where Δ*F* is the energy difference between states A and B, *k_B_* is the Boltzmann constant, *T* is temperature, and *E*_**A**_ is the energy of state A. The angular brackets denote ensemble average over a trajectory for state A.

λ is varied in 32 windows between WT (λ = 0) and mutated (λ = 1) states to maintain overlaps of probability distributions (63). Each window is 200 ps long with 50 ps equilibration and 150 ps data generation. The FEP calculation for the full-length systems are repeated 8 times, with the starting structures selected from the 50, 100, 150 and 200 ns time point of each of the duplicate MD simulations. For N-termini truncated systems, the last snapshots of the 50 ns-long MD simulation for the WT^Δ^_L1_ and WT^Δ^_L2_ complexes are used for the single mutations (WT^Δ^ → G330T^Δ^ and WT^Δ^ → H372A^Δ^), replicated four times. Exponential averaging is used to obtain the reported Δ*G* values.

FEP calculations are compared to the experimental findings as schematically displayed in Figure 2 a. *K_d_* values obtained from binding affinity experiments (23) are employed to calculate standard binding free energies through Δ*G* = −*k_B_T* ln(*K_d_*).

By utilizing the Δ*G* results from the experimental data from Cycle *A* in Figure 2a,

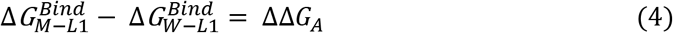

with the FEP results,

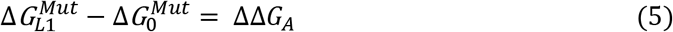

Equations 4 and 5 are arranged as,

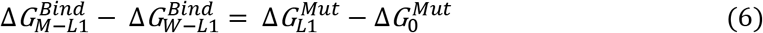

All variables are illustrated in Figure 2 a, and the same calculations are conducted for Cycle *B*. ΔΔΔ*G* is utilized for the validation of the FEP results.

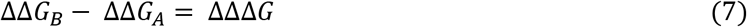

## Acknowledgments

We thank Deniz Sezer for fruitful discussions and the Scientific and Technological Research Council of Turkey (Grant Number 117F389) for funding.

## Supplementary Information

**Figure S1.**
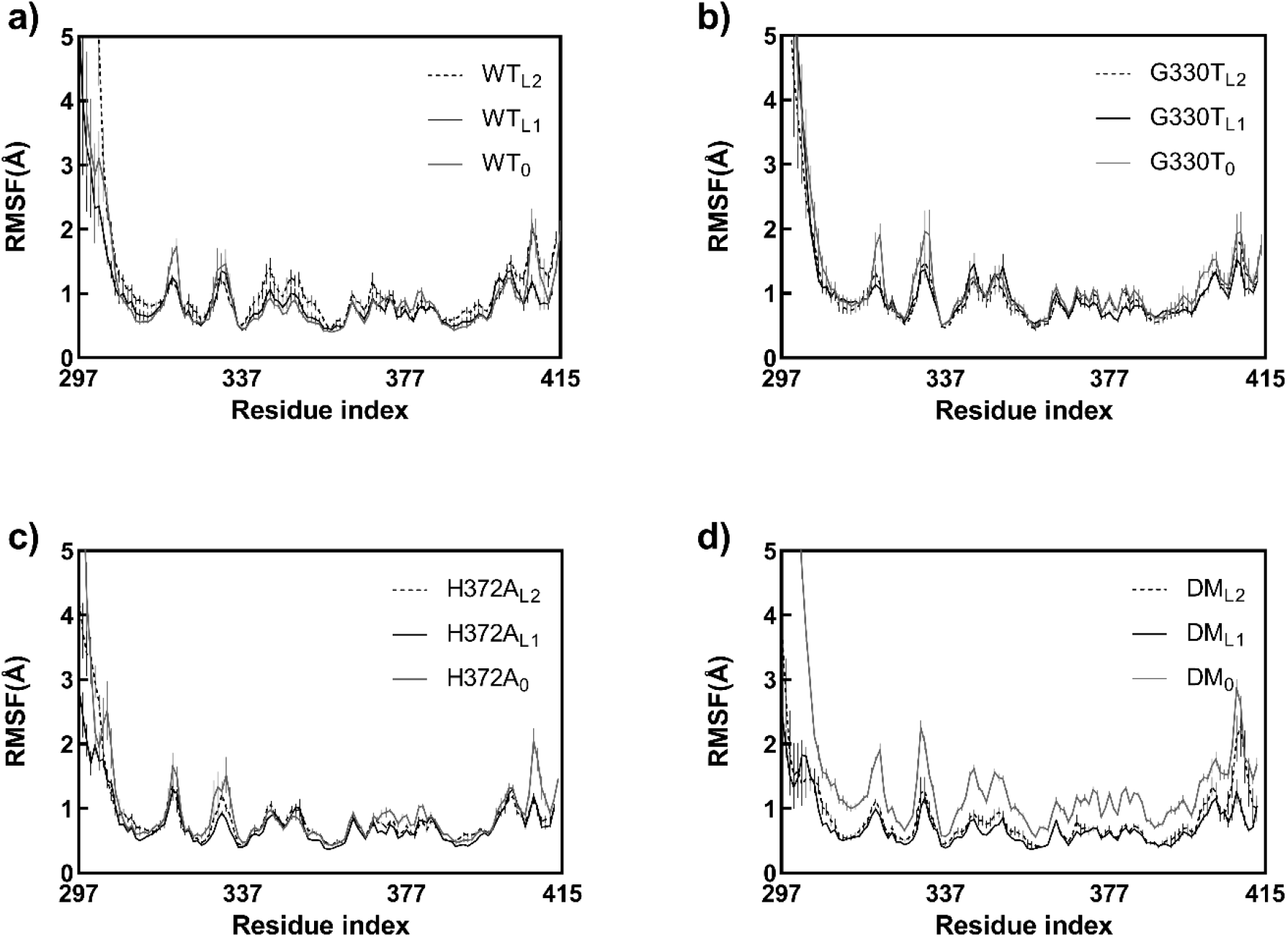
The RMSF results of WT, G330T, H372A and DM for apo and L_1_/L_2_ bound cases. 40 ns-long chunks from duplicate trajectories are averaged; error bars indicate the standard error of these eight chunks. **a, b** WT and G330T show similar patterns and a highly mobile N-terminus region in all cases. **c, d** H372A and DM display similar regimes with a mobile whisker and peak around residue 408. Interestingly, DM_0_ has a significantly high mobility compared to the ligand-bound complexes.

**Figure S2.**
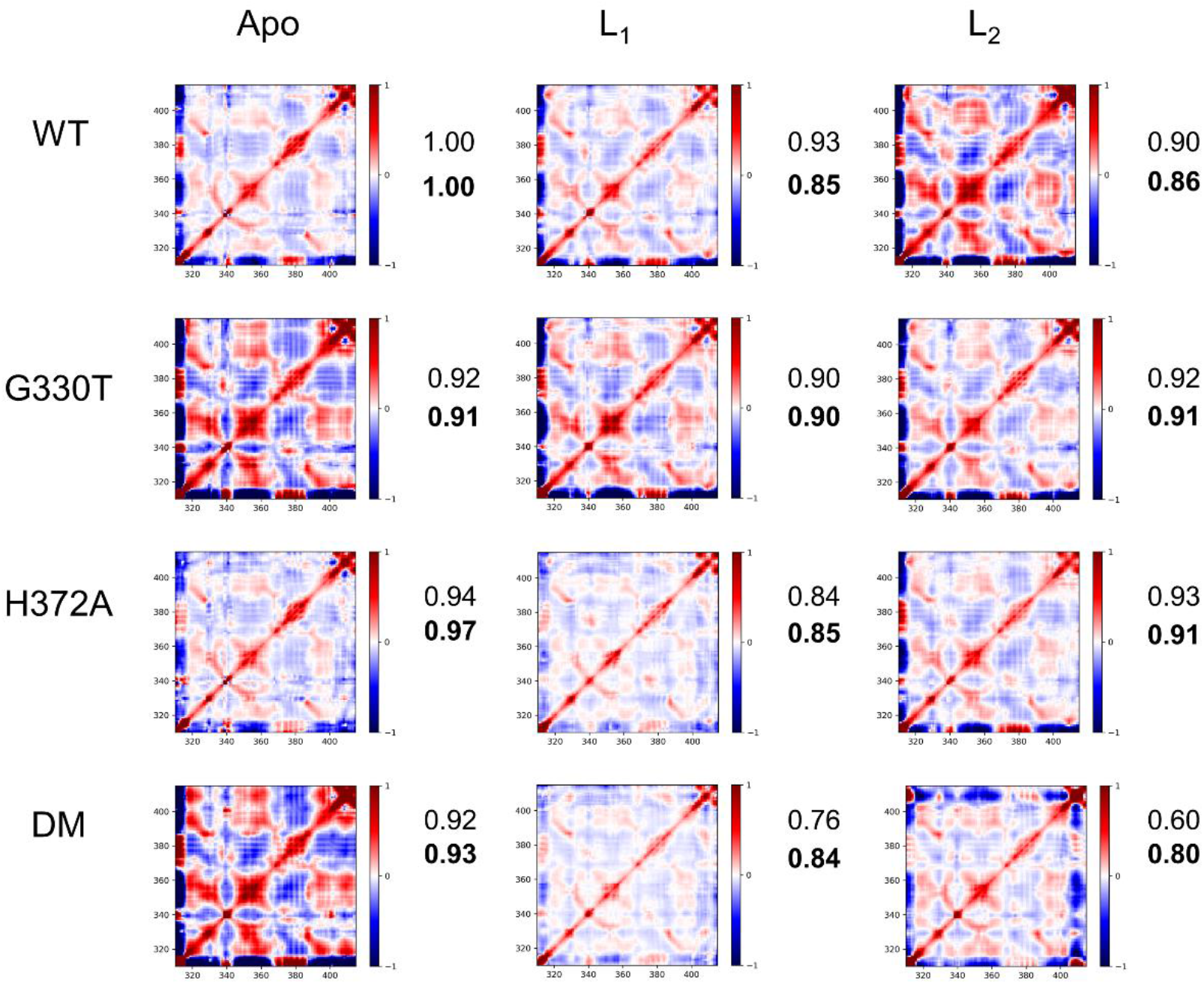
Cross-correlation maps of WT, G330T, H372A and DM for apo and L_1_/L_2_ bound cases. 40 ns-long chunks from duplicate trajectories are averaged for each complex. Graphs are built for the whole protein. Numbers display the similarity to WT_0_, 1 being for identical correlation maps; bold values are computed omitting the N-terminus in the correlation map calculations. DM_L1_ and DM_L2_ display the largest departure in fluctuation patterns compared to the apo WT complex.

**Figure S3.**
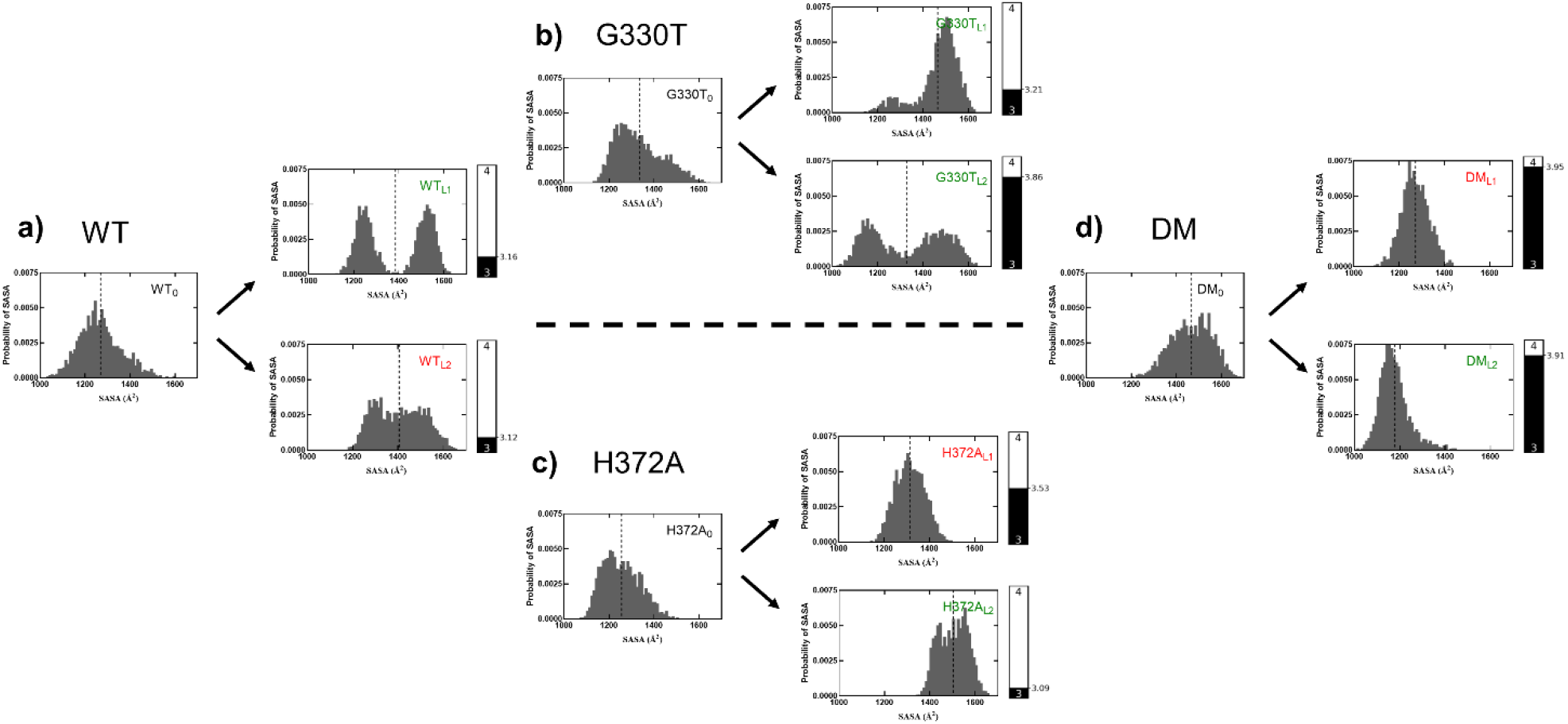
Probability distribution of SASA for the full-length N-terminus; vertical dashed lines indicate average SASA; side bars display the average hydrogen bond count between the PDZ domain and the ligand in the MD simulations in ligand-bound cases. Complexes with favorable binding to the ligands are labelled in green, and complexes with unfavorable binding are in red. These distributions quantify the N-terminus conformations displayed in Figure 3.

**Figure S4.**
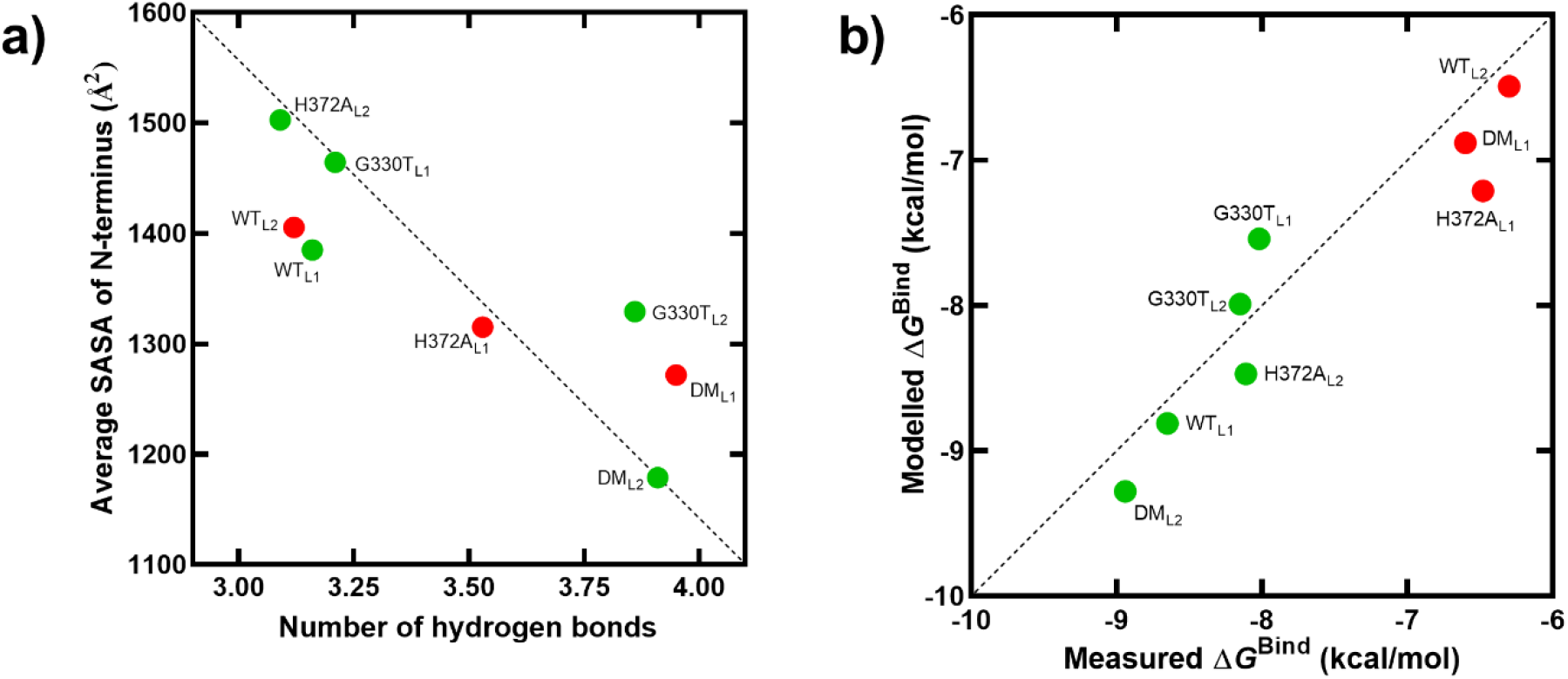
**a** Regression between *N*_H-bonds_ (Table S1) and SASA of N-terminus residues (averages in Figure S3); Pearson *R* = −0.86, *p*-value < 0.01); **b** Regression between modelled (Equation 2) and modelled binding free energies (Table 2); Pearson *R* = 0.94, *p*-value < 0.001) The functional complexes are shown in green, while unfavorable ones are displayed in red.

**Figure S5.**
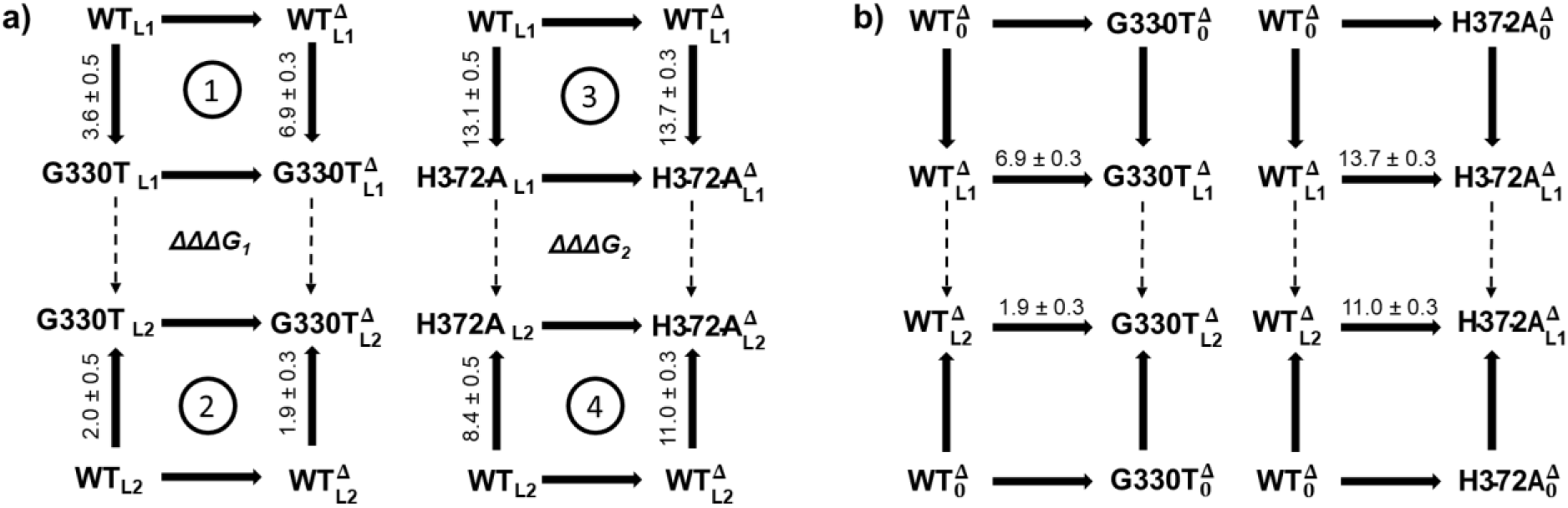
**a** Thermodynamic cycle depicting the role of N-terminus removal on the single mutations. Vertical arrows indicate the mutation process, and the horizontal arrows display the deletion operation on complexes denoted by the superscript Δ. The difference between the horizontal changes are equivalent to that of the vertical ones in each cycle. ΔΔ*G* values of the cycles ① and ④ indicate that the G330T_L1_ and H372A_L2_ forms are highly unfavorable after the operation, while, those for ② and ③ have low or no cost. **b** Thermodynamic cycles depicting mutation and ligand binding of the truncated complexes; ΔΔΔG represent the grand difference between the binding affinities of the WT versus mutant towards either ligand as ΔG of the apo complexes cancel out. 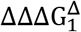 and 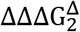 are −5.0 ± 0.4 and −2.7 ± 0.4 kcal/mol, respectively.

**Table S1.** Average number of hydrogen bonds (*N*_H-bonds_) between the protein and the ligand for the full and N-terminus truncated protein structures.

| | Full length protein | Truncated ($\Delta$) protein |
| --- | --- | --- |
| WT<sub>L1</sub> | 3.2 ± 1.4 | 3.4 ± 1.6 |
| WT<sub>L2</sub> | 3.1 ± 1.6 | 3.4 ± 1.7 |
| G330T<sub>L1</sub> | 3.2 ± 1.6 | 3.7 ± 1.7 |
| G330T<sub>L2</sub> | 3.9 ± 1.7 | 4.3 ± 1.7 |
| H372A<sub>L1</sub> | 3.5 ± 1.7 | 3.0 ± 1.4 |
| H372A<sub>L2</sub> | 3.1 ± 1.5 | 3.5 ± 1.7 |
| DM<sub>L1</sub> | 4.0 ± 1.7 | 4.4 ± 1.9 |
| DM<sub>L2</sub> | 3.9 ± 1.7 | 4.3 ± 2.0 |

**Table S2.** Average SASA of the whole protein (*S*) and SASA variances of N-terminus region for charged residues (*σ*) used in Equation 2.

| | $S$ | $\sigma$ |
| --- | --- | --- |
| WT<sub>0</sub> | 7128 | 60 |
| WT<sub>L1</sub> | 6746 | 32 |
| WT<sub>L2</sub> | 6736 | 51 |
| G330T<sub>0</sub> | 7171 | 58 |
| G330T<sub>L1</sub> | 6875 | 40 |
| G330T<sub>L2</sub> | 6818 | 47 |
| H372A<sub>0</sub> | 7105 | 66 |
| H372A<sub>L1</sub> | 6734 | 50 |
| H372A<sub>L2</sub> | 6937 | 31 |
| DM<sub>0</sub> | 7300 | 62 |
| DM<sub>L1</sub> | 6533 | 81 |
| DM<sub>L2</sub> | 6506 | 44 |

